# RIOT - Rapid Immunoglobulin Overview Tool - annotation of nucleotide and amino acid immunoglobulin sequences using an open germline database

**DOI:** 10.1101/2024.08.12.607568

**Authors:** Paweł Dudzic, Bartosz Janusz, Tadeusz Satława, Dawid Chomicz, Tomasz Gawłowski, Rafał Grabowski, Przemysław Jóźwiak, Mateusz Tarkowski, Maciej Mycielski, Sonia Wróbel, Konrad Krawczyk

## Abstract

Antibodies are a cornerstone of the immune system, playing a pivotal role in identifying and neutralizing infections caused by bacteria, viruses, and other pathogens. Understanding their structure, and function, can provide insights into both the body’s natural defenses and the principles behind many therapeutic interventions, including vaccines and antibody-based drugs. The analysis and annotation of antibody sequences, including the identification of variable, diversity, joining, and constant genes, as well as the delineation of framework regions and complementarity-determining regions, is essential for understanding their structure and function. Currently analyzing large volumes of antibody sequences is routine in antibody discovery, requiring fast and accurate tools. While there are existing tools designed for the annotation and numbering of antibody sequences, they often have limitations such as being restricted to either nucleotide or amino acid sequences, reliance on non-uniform germline databases, or slow execution times. Here we present Rapid Immunoglobulin Overview Tool (RIOT), a novel open-source solution for antibody numbering that addresses these shortcomings. RIOT handles nucleotide and amino acid sequence processing, comes with a free germline database, and is computationally efficient. We hope the tool will facilitate rapid annotation of antibody sequencing outputs for the benefit of understanding antibody biology and discovering novel therapeutics.

**Availability:** RIOT is available at https://github.com/NaturalAntibody/riot_na.

## Introduction

Antibody diversity and specificity are crucial for the immune system’s ability to recognize and combat a wide range of pathogens, leading to effective acquired immunity. Humans can produce a multitude of unique antibodies through V(D)J recombination, a process specific to B-cell gene segments (Chi, Li, and Qiu 2020). Somatic hypermutation further diversifies these antibodies, optimizing their affinity for antigens (Wagner and Neuberger 1996). Antibodies are highly specific due to the unique antigen-binding sites formed by the variable regions. This specificity is exploited in monoclonal antibodies (Crescioli et al. 2024), which are identical antibodies from a single B cell clone used in medical treatments for diseases such as cancer or autoimmune disorders. The complexity and diversity of antibody repertoires, underpinned by the mechanisms of V(D)J recombination and somatic hypermutation, necessitate sophisticated tools for their analysis and interpretation.

Antibody numbering schemes, such as the Kabat (Kabat 1987), Chothia (Al-Lazikani, Lesk, and Chothia 1997), Martin (Abhinandan and Martin 2008) and IMGT (Lefranc et al. 2003) and others (Honegger and Plückthun 2001; Patel et al. 2023), provide a standardized framework for annotating the amino acid positions in antibody sequences. This standardization is essential for comparing antibodies, correct delineation of Complementarity-determining regions (CDRs), humanization, understanding their structure-function relationships, and multiple other applications. The numbering schemes are not only fundamental for academic research but also play a crucial role in the development of monoclonal antibodies for therapeutic use.

The widely recognized methods for antibody numbering predominantly fall into two categories: those that rely on sequence alignment in the traditional sense, and those based on Hidden Markov Models (HMM) alignments (Eddy 2011). Both approaches necessitate a substantial repository of previously annotated antibody sequences to construct the necessary reference databases. Sequence-based methods can perform alignments of input sequence either to pre-annotated antibodies sequences or to germline genes. While the reliance on pre-annotated sequences yields effective results for recognized patterns, it encounters limitations with atypical ones.

An example of profile-based numbering, which applies higher significance to conserved regions, was introduced in AbNum (Abhinandan and Martin 2008). Here, authors assign framework-derived profile segments to input sequence segments. Those are then used to determine region boundaries and finally region sequences are aligned to consensus patterns. While the authors thoroughly investigated the correctness of the numbering results, the method does not provide germline assignment.

Sequence-alignment-based numbering methods offer simplicity and accuracy for well-represented sequences but require a good-quality input germline database. For example, IgBLAST (Ye et al. 2013) which aligns input sequences to germline genes to determine regions in the input sequence, achieves good germline assignment accuracy, yet is computationally expensive and has a limited number of supported numbering schemes. In contrast, HMM-based methods provide robustness and adaptability for unseen sequences at the cost of even higher complexity and computational expense. To generate HMM profiles used as a reference database, scheme-gapped pairs of V and J genes can be used as it was implemented in ANARCI (Dunbar and Deane 2016). This approach however yields alignment penalization of unusually long CDRs.

Another sequence-based numbering that addressed the robustness problem was implemented in AbRSA (Li et al. 2019). In their approach, authors prepared numbered antibody consensus sequences of heavy and light chains, where each position in the consensus sequence is a list of the most popular antibody residues constructed from the database abYsis (Swindells et al. 2017). The query sequence is aligned to consensus sequence using a modified Needleman-Wunch algorithm which considers consensus residue region when calculating the alignment score. While this approach correctly assigns lower significance to hypervariable regions, it is limited with the number of available numbering schemes and it does not provide germline assignment.

Further speed improvement with respect to AbRSA was taken by AntPack (Parkinson and Wang 2024). The method is the fastest currently, however it only handles amino acids and germline genes are assigned using sequence identity rather than obtained by more accurate alignments.

The available tools either focus purely on annotation of amino acids or nucleotide sequences and are dependent on the underlying database. IMGT and efforts by the AIRR community continue to be the main sources of germline genes (Lefranc 2003; W. Lees et al. 2020; W. D. Lees et al. 2023; Omer et al. 2020). Nonetheless, these undergo frequent updates which can lead to inconsistencies between annotations employing a particular version of each database. Employing different pieces of software for annotation and different germlines results in inconsistent annotations (Smakaj et al. 2020). Standardization of the annotation protocols across nucleotides/amino acids, as well as the underlying germline database, would increase consistencies between analyses.

Here, we present RIOT, software integrating nucleotide and amino acids immunoglobulin sequence annotation with built-in human and mouse germline databases. We benchmark RIOT against leading tools in nucleotide-sequence annotation and amino-acid sequence annotation showing a best in class versatility and a significant decrease in annotation time. We hope that our software will help in the analysis of antibodies by offering a single solution for sequence analysis with a standardized reference germline database.

## Methods

### Annotation pipeline

Here we present RIOT - Rapid Immunoglobulin Overview Tool - sequence-based antibody numbering software that facilitates fast execution times, accurate germline assignments, and versatile selection of numbering schemes.

At first, V gene candidates are selected in k-mer matching-based prefiltering process and the top 12 matching genes are pairwise aligned using the Striped Smith-Waterman algorithm (Farrar 2007) to the input sequence in forward and reverse complemented directions. The gene with the best e-value is assigned to the sequence together with source locus and organism. Then the part of the input sequence that was aligned is masked so the J gene can be identified on the remaining part of the sequence. Only J genes that match the assigned organism and locus are considered. After further masking the part of the sequence which is aligned to the J gene, the remaining parts of the sequence are used to identify C gene and in the case of heavy chains - D gene (Figure 1). For nucleotide sequences on the input, the reading frame is inferred from V gene alignment so the input sequence is translated to amino acids sequence which are used to align the input sequence to the scheme of choice. After numbering, additional validations are being performed.

**Figure 1.**
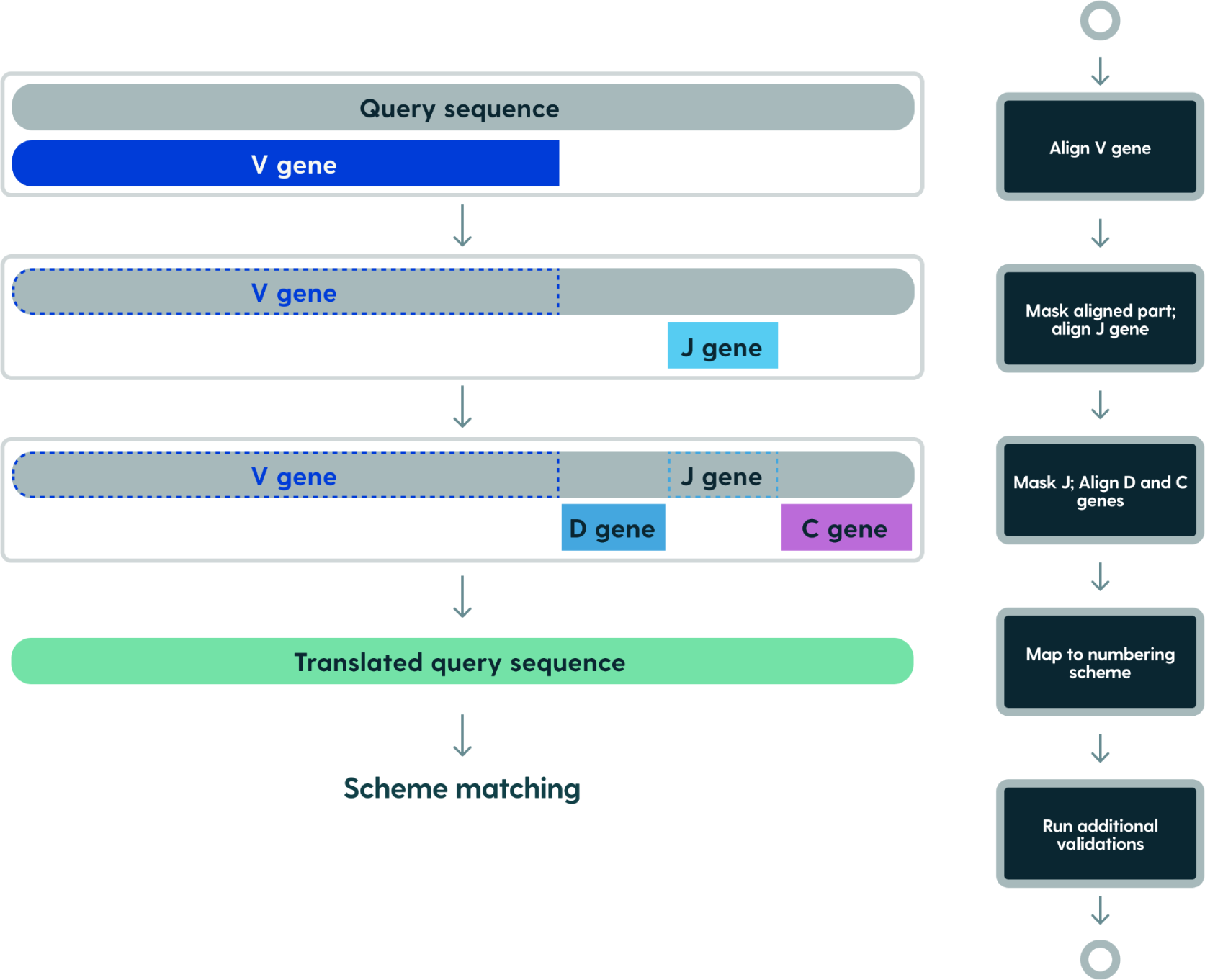
Annotation pipeline. Align the query sequence to V genes (both forward and reverse complemented). Infer the organism, locus, and reading frame from the best V gene alignment. Mask the query sequence aligned with the V gene. Align the J gene, offsetting the best alignment by the masked V region. Mask again the query sequence, and align D and C genes. Infer reading frame from V gene alignment. Translate amino acid alignments to a selected numbering scheme (Kabat, Chothia, Martin, or IMGT). The pipeline is identical for nucleotides and amino acids with the exception of a lack of D/C gene annotation in amino acids.

### Scheme mapping / numbering

In contrast to the majority of numbering tools, RIOT does not align query sequences to full-length scheme-gapped sequences or profiles. Rather it aligns V and J segments of the query sequence separately, to ungapped (non scheme aligned) germline sequences. Query-gene alignments are then merged with already known gene-scheme alignments to produce query-scheme mapping on corresponding segments. This approach renders reliable germline assignments and avoids penalization of unusually long CDRs which is common in artificial sequences deposited in patents (Krawczyk, Buchanan, and Marcatili 2021) (Figure 2).

**Figure 2.**
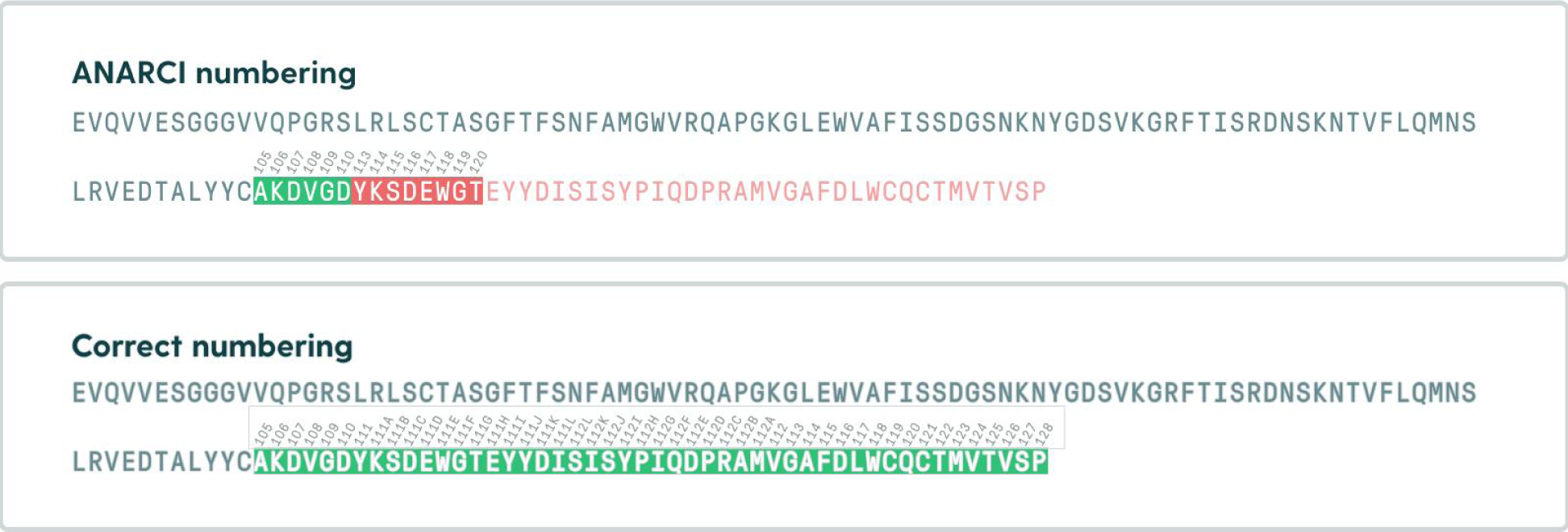
Example of an issue with alignment to artificially gapped sequences. Alignment to IMGT gapped profiles created from V-J gene pairs leads to penalization of unusually long CDR (PDB: 4ocr). To address this RIOT aligns V and J genes separately. The example above was generated using ANARCI adapted from (Robinson 2023).

We argue that alignment to ungapped genes sequences will yield more accurate germline assignments as we are aligning input to biologically accurate genes rather than sequences where artificially introduced gaps can change alignment scores because of the gap penalties. As a consequence of this, a good quality germline database is needed on input to provide reliable numbering.

We know how gene amino acids sequences map to schemes therefore we are able to efficiently combine query-gene alignment with gene-scheme alignment to align input to the scheme of choice. This mapping process is performed by the sequence of steps given in Figure 3.

**Figure 3:**
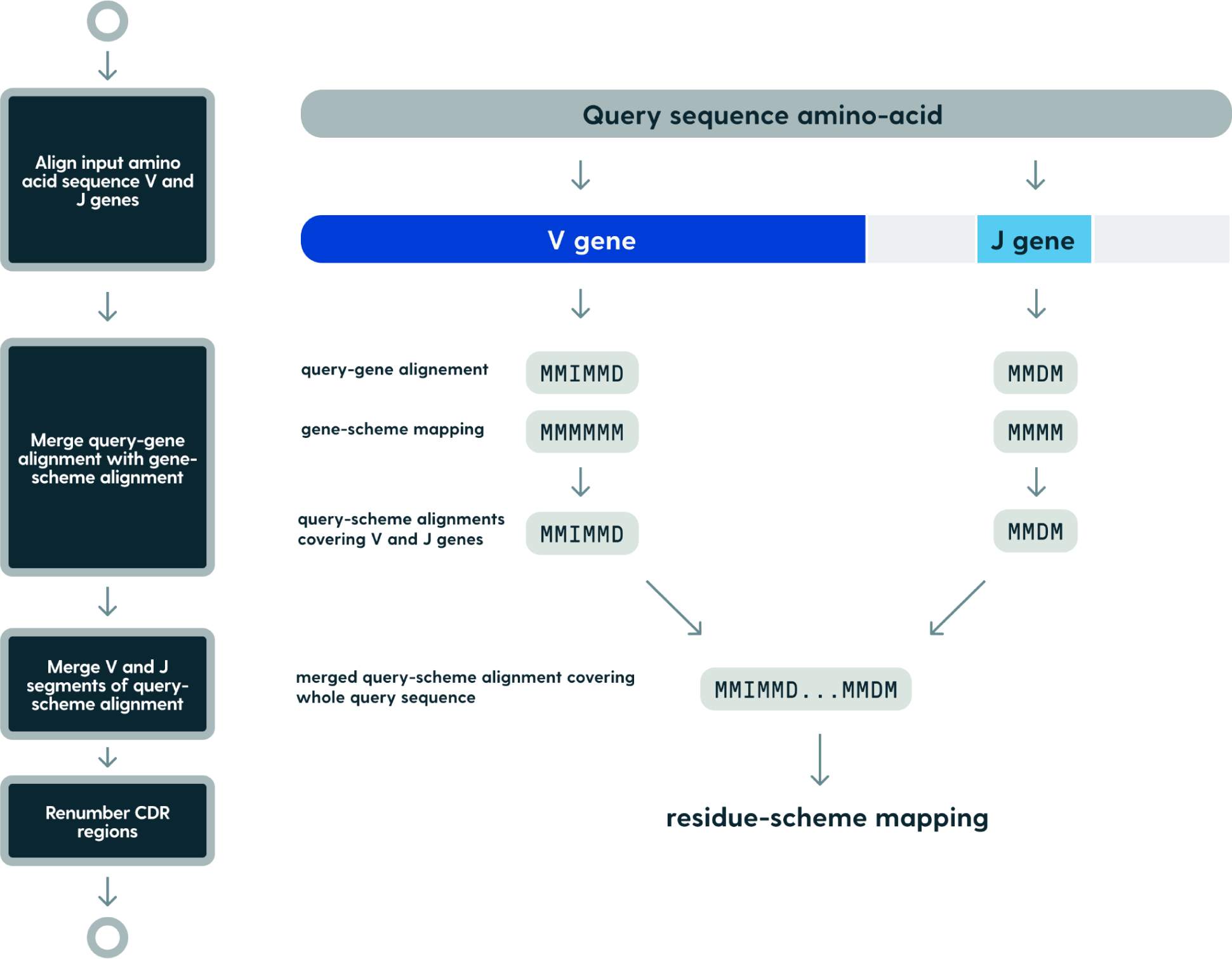
Applying numbering schemes. Query amino acid sequence is aligned to germlines to obtain query-gene alignment. This mapping is merged with known gene-scheme mapping for scheme of choice. This way query-scheme mappings are obtained for V and J segments which are subsequently joined to create query-scheme mapping covering the whole input sequence.

Input amino acid sequence is aligned to amino acid sequences of V and J genes. In the case of nucleotide pipeline prefiltering step is not executed as the genes from best nucleotide alignments are used. Then, query-gene alignments are merged with gene-scheme alignments to produce query-scheme mapping for V and J derived fragments of input sequence. The merging is executed by following query-gene and gene-scheme alignment backtraces. This way produced query-scheme alignments for V and J segments are then collapsed so that all consecutive insertions and deletions are converted to matches. In the next step, V and J query-scheme alignments are merged together so full query-target scheme alignment is generated which allows to determine regions boundaries. To infer the scheme mapping between V and J alignments, the difference between scheme legal positions and the length of the actual unaligned query sequence segment is calculated. If the query fragment between V and J alignments is longer than the number of legal scheme positions, the segment alignment consists of the number of matches that are allowed in the scheme with additional insertions put in the middle. Similarly, if the query sequence is shorter, the middle segment consists of matches in count equal to the length of the sequence and missing scheme positions are filled as deletions in the middle. Finally, CDR regions are renumbered so that they match scheme definitions - insertions are put after the specified position (or in the middle for IMGT scheme) and deletions are put before the specified position. If there are too many deletions to be put before the specified position, they are appended after the specified place.

Framework and CDRs regions boundaries are inferred from query sequence - scheme alignment using the definitions provided by Andrew Martin (Table 1) (http://www.bioinf.org.uk/abs/info.html).

**Table 1:**
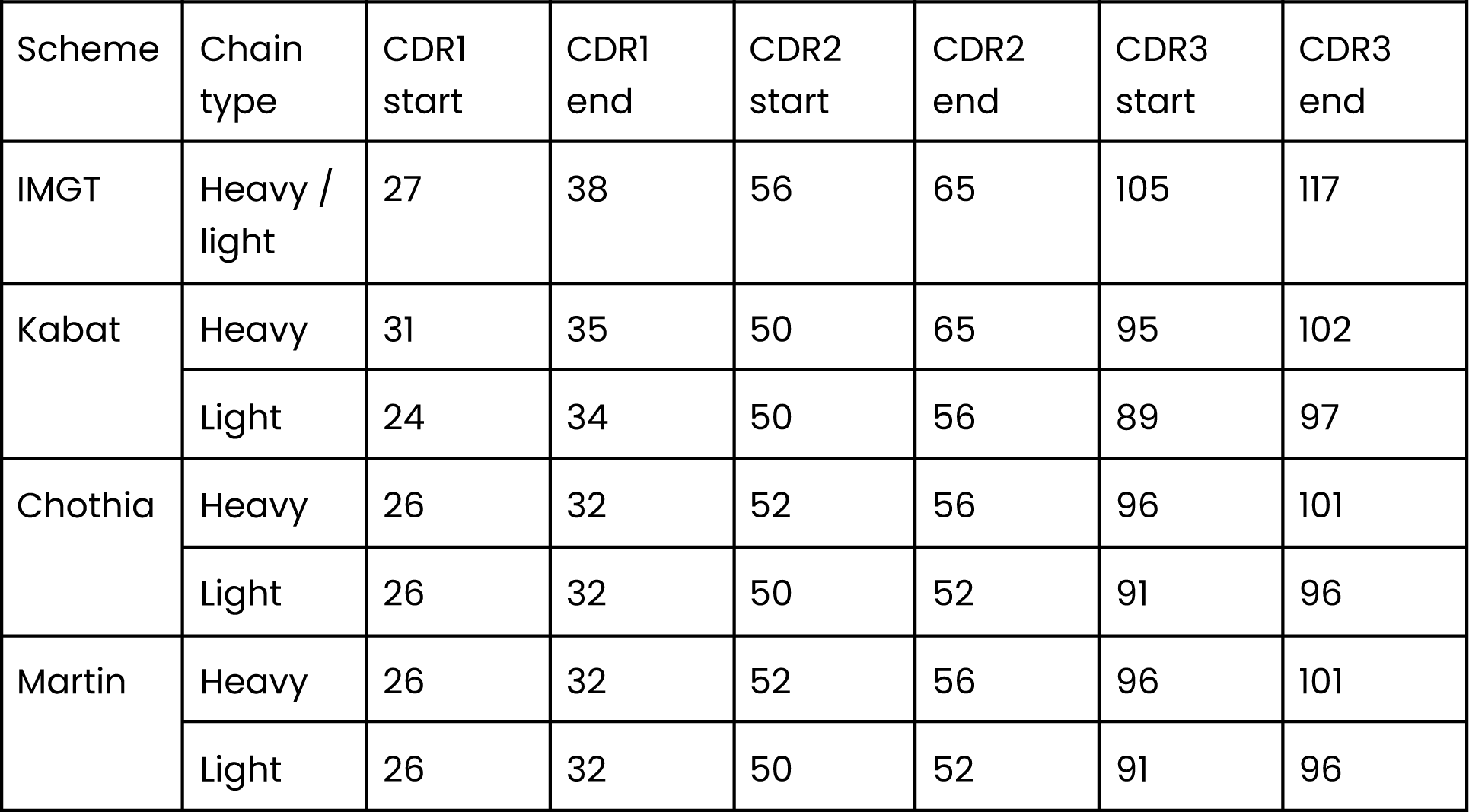
CDR regions boundaries according to each numbering scheme.

| Scheme | Chain type | CDR1 start | CDR1 end | CDR2 start | CDR2 end | CDR3 start | CDR3 end |
| --- | --- | --- | --- | --- | --- | --- | --- |
| IMGT | Heavy / light | 27 | 38 | 56 | 65 | 105 | 117 |
| Kabat | Heavy | 31 | 35 | 50 | 65 | 95 | 102 |
|  | Light | 24 | 34 | 50 | 56 | 89 | 97 |
| Chothia | Heavy | 26 | 32 | 52 | 56 | 96 | 101 |
|  | Light | 26 | 32 | 50 | 52 | 91 | 96 |
| Martin | Heavy | 26 | 32 | 52 | 56 | 96 | 101 |
|  | Light | 26 | 32 | 50 | 52 | 91 | 96 |

Kabat, Chothia and Martin schemes perform numbering in a distinct fashion to IMGT by employing indel positions. The indel positions are anchors for insertions and deletions. Insertions are placed after the specified indel residue position and deletions before. Anchor positions for all CDRs are defined in Table 2.

**Table 2:** CDR indel positions for Kabat-derived schemes.

| Scheme | Chain type | CDR1 | CDR2 | CDR3 |
| --- | --- | --- | --- | --- |
| Kabat | Heavy | 35 | 52 | 100 |
|  | Light | 27 | 52 | 95 |
| Chothia | Heavy | 31 | 52 | 100 |
|  | Light | 30 | 52 | 95 |
| Martin | Heavy | 31 | 52 | 100 |
|  | Light | 30 | 52 | 95 |

While preparing scheme mappings we noticed that some genes do not comply with scheme definitions by having additional insertion places not defined by schemes. We decided to add these insertions; otherwise such positions would cause misalignment of germlines sequences. This behavior is consistent with the other numbering tools. We numbered the V genes sequences with novel insertions with ANARCI and AbRSA and both programs added insertions in places not allowed by schemes. Insertion places inferred from germline sequences are defined in Table 3.

**Table 3:** Novel germline-inferred insertion positions in Kabat-like numbering schemes.

| Chain type | Position | Schemes | Example |
| --- | --- | --- | --- |
| Light | 66 | Kabat, Chothia | IGLV6-57*01, IGLV5-39*01, IGLV5-45*01 |
| Light | 52 | Kabat, Chothia | IGLV5-45*01, IGLV4-60*03 |
| Light | 68 | Martin | IGLV6-57*03, IGLV5-39*01 |

### Gene matching

Gene matching process consists of two steps: prefiltering used for candidate gene selection and pairwise alignments of the selected gene candidates against query sequence. Candidate with the highest e-value is assigned as the matching gene.

In 2017 Martin Steingger and Johannes Soedig demonstrated significant speed improvements in sequence search over BLAST (Altschul et al. 1990) by introducing diagonal based “prefiltering” step before the alignment in their Mmseqs2 pipeline (Steinegger and Söding 2017). In this work we implemented this concept for the purpose of fast V, D, J and C genes matching.

Prefiltering is a crucial step in the RIOT gene matching pipeline that helps reduce the computational time and resources required for the alignment process. In this step, an index table for target sequences is created. For each k-mer occurring in the target gene sequence, one can look up the target’s identifiers and k-mer positions. These k-mers are then used to identify the best matching gene candidates to the query sequences by calculating diagonal scores. If we construct a coordinate plane in which we denote the x-axis as the position in the query sequence and the y-axis as the position in the target gene, all k-mer matches between the input sequence and target gene can be represented as short diagonal segments starting at coordinates (query position, target position). Consecutive segments can be directly connected to form contiguous sequences of consecutive matching positions. If there is an insertion in the query sequence with respect to the gene sequence, it will be represented as a segment shifted along the x-axis. Similarly, a deletion will be represented as a shift along the y-axis. Segments with a difference between x and y positions in consecutive matches greater than the threshold are discarded. Finally, the number of matching residues serves as a coverage (or diagonal) score, which is used further to select the best matching candidates (Figure 4). By setting a threshold on this score, RIOT filters out sequence pairs that are unlikely to share significant similarity, retaining only the most promising candidates for further processing. This prefiltering step dramatically reduces the number of sequence pairs that need to be considered in the subsequent, more computationally intensive pairwise alignment.

**Figure 4:**
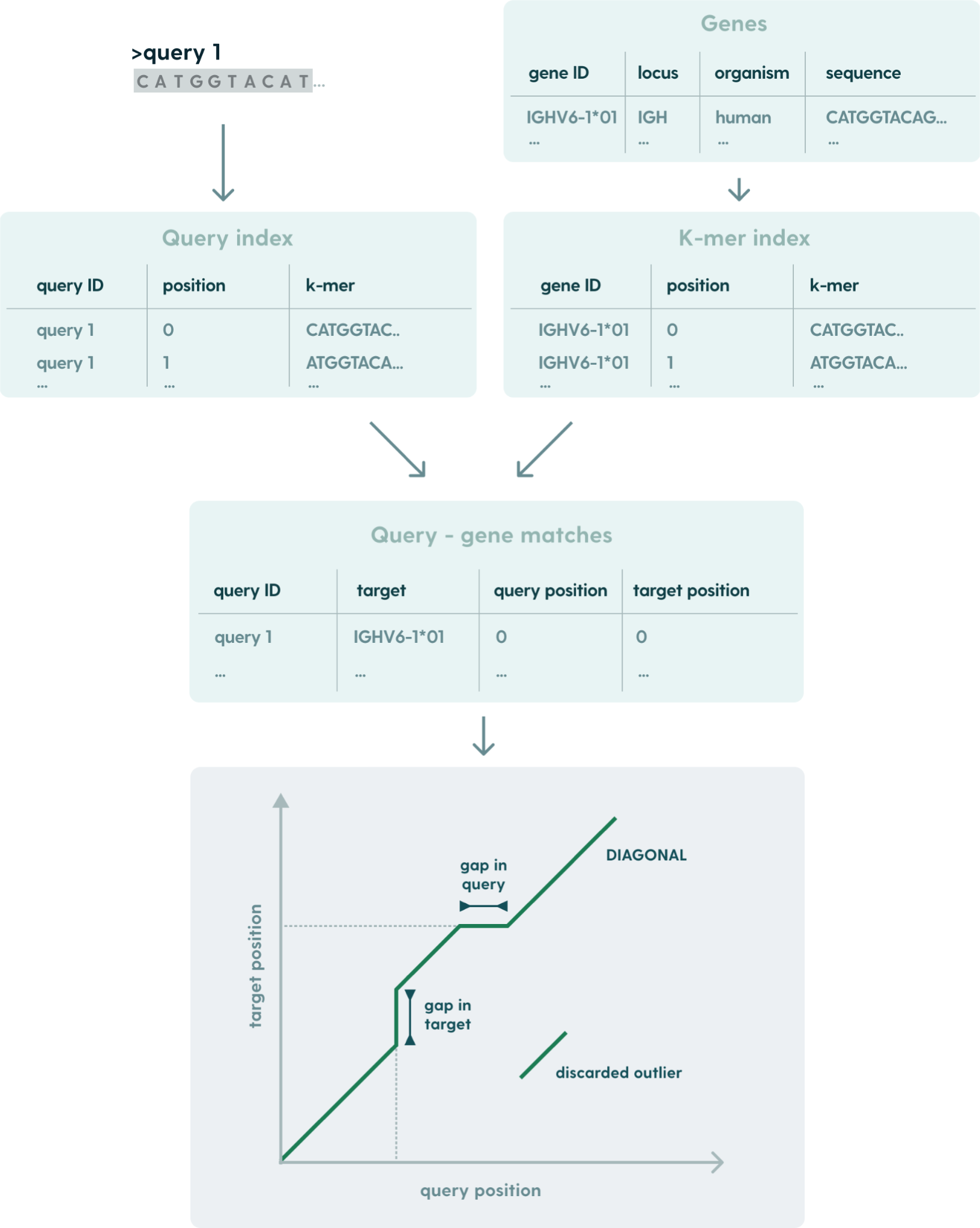
Prefiltering: diagonal score-based candidate genes selection. Representation of diagonal score calculated for single pair of input query and gene.

Prefiltering can be fine-tuned on the number of best matches returned, size of k-mers for the index, distance threshold and sampling threshold. While the sampling threshold and k-mer size impacts the number of k-mer comparisons, which has a direct influence on prefiltering performance, when it comes to the number of best matches returned - prefiltering execution time does not change itself but this alters the number of pairwise alignments performed downstream. As consecutive k-mer matches constitute diagonal construction, distance threshold is the maximum relative offset between query and target positions for k-mer to be included in the diagonal. Finally, the sampling threshold determines the offset between consecutive k-mers from the query sequence.

RIOT prefiltering parameters were fine-tuned using grid search for each gene with the goal of minimizing the number of consecutive alignments while preserving germline alignment accuracy on sample dataset of human sequences. All RIOT prefiltering parameters used are summarized in Table 4.

**Table 4:** Prefiltering parameters for genes. Top N is the number of sequences passing prefiltering to subsequent pairwise alignments. K-mer size determines the length of k-mers used in the prefiltering index. Distance threshold is the maximum relative offset in query and target positions between consecutive k-mer matches. Modulo N is the sampling threshold determined by offset between consecutive k-mers in the query sequence.

| Gene | Locus | Top N | K-mer size | Distance threshold | Modulo N |
| --- | --- | --- | --- | --- | --- |
| V | all | 12 | 9 | 13 | 2 |
| D | IGH | 5 | 5 | 3 | 1 |
| J | IGH | 5 | 5 | 3 | 2 |
| J | IGK | 5 | 5 | 7 | 2 |
| J | IGL | 5 | 5 | 5 | 1 |
| C | IGH | 5 | 5 | 5 | 2 |
| C | IGK | 1 | 5 | 3 | 1 |
| C | IGL | 5 | 3 | 5 | 1 |

To further speedup the gene candidates selection, we used the *ahash* library (https://docs.rs/ahash/latest/ahash/) for rapid hash calculations of k-mers in index lookups.

After the prefiltering step in the RIOT gene matching pipeline, the remaining candidate sequence pairs proceed to the pairwise alignment stage. This step employs the SIMD optimized Smith-Waterman algorithm (Farrar 2007) for the selected query-target sequence pairs. The alignment information is stored as a CIGAR (Compact Idiosyncratic Gapped Alignment Report) string, which is a concise representation of the consecutive mapping operations (match, mismatch, insertion, and deletion) and their counts. Finally, alignment scores are calculated, such as sequence identity or e-value. Resulting alignments are sorted by e-value, so it is easy to pick the best one in the last step. Alignment parameters used in RIOT are the same as defaults in IgBLAST. We found them to produce accurate alignments and at the same time similar to what the user base is accustomed to. They are summarized in Table 5.

**Table 5:** Pairwise alignment parameters.

| Sequence type | Score | Gap open penalty | Gap extend penalty |
| --- | --- | --- | --- |
| Nucleotides | Match: 1; Mismatch: -1 | 4 | 1 |
| Amino acids | BLOSUM62 | 11 | 1 |

### Germline Database

The Open Germline Receptor Database (OGRDB) (W. Lees et al. 2020) is used as a primary source of germline genes in RIOT. It is a reference database for inferred immune receptor genes which addresses the need for comprehensive germline gene reference sets to interpret AIRR-seq data accurately. Current reference sets are incomplete and under-representative of human and animal population diversity (Peng et al. 2021). OGRDB provides V, D and J genes sequences for human and mouse with supporting evidence. Mouse sequences from all strains were put to a single database and deduplicated. Human C genes were obtained from NCBI (Maglott et al. 2005) and are reused for all species. Allele abundance for gene sequences is presented in Table 6.

**Table 6.** OGRDB-derived internal germline database in RIOT. Allele abundance is given for each locus in human and mouse databases.

| Organism | Segment | H | K | L |
| --- | --- | --- | --- | --- |
| Human | V | 190 | 62 | 75 |
|  | D | 31 | - | - |
|  | J | 7 | 7 | 9 |
|  | C | 9 | 1 | 5 |
| Mouse | V | 643 | 459 | 7 |
|  | D | 19 | - | - |
|  | J | 6 | 11 | 7 |

### Availability

RIOT is open-sourced via the following repository: https://github.com/NaturalAntibody/riot_na and is free to use for non-commercial use by non-commercial organizations.

## Results

### Feature comparisons of existing tools

We compared RIOT to other tools in two dimensions: supported numbering schemes (Table 7) and annotation possibilities (Table 8).

**Table 7.** Comparison of supported numbering schemes in existing tools. *IgBlast does not provide residue-level annotations like all other tools.

|  | AntPack | AbNum | AbRSA | ANARCI | RIOT | IgBLAST |
| --- | --- | --- | --- | --- | --- | --- |
| Kabat | yes | yes | yes | yes | yes | yes* |
| Chothia | no | yes | yes | yes | yes | no |
| Martin | yes | yes | no | yes | yes | no |
| IMGT | yes | no | yes | yes | yes | yes* |
| Aho | no | no | no | yes | no | no |

**Table 8.** Comparison of supported features of existing antibody numbering tools. *IgBlast only has limited functionality regarding amino acid annotation as it only aligns V-genes.

|  | AntPack | AbNum | AbRSA | ANARCI | RIOT | IgBLAST |
| --- | --- | --- | --- | --- | --- | --- |
| Nucleotide annotation | no | no | no | no | yes | yes |
| Amino acid annotation | yes | yes | yes | yes | yes | yes* |
| Germline assignment | yes | no | no | yes | yes | yes |
| Isotype assignment | no | no | no | no | yes | yes |
| AIRR format output | no | no | no | no | yes | yes |

Considering the supported numbering schemes, RIOT falls behind ANARCI by not supporting the Aho numbering scheme. This was a conscious choice since there are multiple numbering systems, not all listed here and we chose to implement only those that are commonly used within industry and academic community. While IgBLAST supports Kabat and IMGT in its annotations it does not provide residue-level annotations.

Regarding annotation features, we list the ability to handle nucleotide or amino acid formats, germline assignment, isotype assignment, and compatibility with the AIRR format (Table 8). From all presented tools, only RIOT and IgBLAST checks all boxes in compared features. IgBLAST support for amino acids is limited as it is able to align only V genes.

### Germline assignment accuracy and speed

To assess the performance of RIOT, we compared it to the leading annotation tools available: IgBLAST, ANARCI and AntPack. AbNum nor AbRSA were not included in the comparisons as they do not provide germline assignments functionality. We constructed two benchmarking datasets: containing nucleotide and amino acids sequences to benchmark IgBLAST and ANARCI+AntPack respectively. Nucleotides sequence dataset was constructed by downsampling the AbNGS (Dudzic et al. 2024) dataset and amino acids sequence dataset was derived from therapeutics sequences curated from INN entries (Młokosiewicz et al. 2022). All tools tested were configured to use the same germline database: In case of IgBLAST, we provided OGRDB germline database as an input, and for ANARCI we constructed HMM profiles from V-J gene pairs gapped to IMGT. The IMGT-gapped OGRDB germlines were also provided to AntPack for consistency. In all cases, we assessed the accuracy of assigning germlines as well as wall-clock running time.

To construct ground-truth datasets in both cases, each test-set sequence was aligned to all of the germline gene sequences using the Striped Smith-Waterman algorithm, and the gene with the lowest e-value score was assigned as a ground truth. We evaluated the germline assignment accuracy on the level of genes and alleles. As each experiment was performed 10 times on an 8-core machine, execution wall clock time was measured and mean value from 10 distinct runs was used to compare the performance.

IgBLAST achieves comparable gene assignment accuracy to RIOT (Table 9). Our software, however, achieves significant speed improvement over IgBLAST, being more than 4x faster. When working with highly mutated sequences or incomplete germline databases for novel organisms, prefiltering parameters should be further adjusted with fine-tuning.

**Table 9.** Comparison of gene assignment accuracy and execution time for IgBLAST and RIOT. Execution time reported is wall-clock from execution on an 8-core machine.

|  |  | RIOT | IgBLAST | Dataset |
| --- | --- | --- | --- | --- |
| Gene | V | 99.85% | 99.88% | NGS sample stratified by<br>bioproject (362180 sequences)<br>Homo Sapiens |
|  | J | 99.91% | 98.84% |  |
| Allele | V | 99.83% | 99.73% |  |
|  | J | 99.91% | 98.80% |  |
| Execution time 8 cores |  | 00:04:26 | 00:18:02 |  |

The therapeutics-derived dataset was used to measure the germline assignment accuracy for amino acid-based annotation using ANARCI and AntPack. Because of its small size, the execution time measurements were not reliable. Therefore to measure the processing efficiency, we constructed another amino acids dataset by translating NGS sequences. Germline amino acid sequences were deduplicated as there may be multiple distinct alleles with the same translation sequence.

RIOT outperforms ANARCI on all metrics (table 10). As we investigated, surprisingly low germline assignment accuracy of ANARCI could be attributed to its germline assignment method: iterating over the alignment to select the gene with the highest sequence identity. We advocate for assigning genes using e-values from alignments as we believe this renders more biologically meaningful results rather than sequence identity alone. Since AntPack has a similar germline annotation scheme as ANARCI, it is outperformed by RIOT on this count. Nevertheless AntPack achieves a significantly lower run-time of 32 seconds vs 2 minutes and 40 seconds in RIOT.

**Table 10.** Comparison of gene assignment accuracy and execution time for ANARCI, AntPack and RIOT. Execution time reported is wall-clock from execution on an 8-core machine. *AntPack assigns genes only in the IMGT scheme - assignments in other schemes require double numbering. Time measurements show the numbering to the IMGT scheme.

|  |  | RIOT | ANARCI | AntPack | Dataset |
| --- | --- | --- | --- | --- | --- |
| Gene | V | 96.94% | 65.42% | 80.04 | Therapeutics heavy and light separate (1274 sequences)<br>V genes, Homo Sapiens |
|  | J | 97.72% | 78.94% | 77.99% |  |
| Allele | V | 96.86% | 62.17% | 76.35% |  |
|  | J | 97.65% | 78.60% | 77.77% |  |
| Execution time 8 cores |  | 00:02:40 | 00:18:01 | 00:00:32 | 362180 random sequences from AbNGS |

### Numbering accuracy

In order to evaluate the numbering accuracy, therapeutics amino acids sequence dataset was used. All sequences were numbered with AbNum, AbRSA, ANARCI, AntPack and RIOT to numbering schemes given in Table 7. Numberings were converted to a common format so that all numbering results have unified insertion notation. Finally, the results from all tools were aligned together so non-consensus sequences could be identified. AbRSA numbering results were excluded from comparisons as it caused most conflicts (CDR numbering does not comply with scheme definitions).

### IMGT - ANARCI, AntPack and RIOT comparison

Out of all therapeutics derived variable regions, there were only 23 cases where tools rendered conflicting numberings. Upon examination of the conflicting numberings, we observed that some of the differences can be attributed to different CDR smoothing heuristics between programs. Additionally, the numbering differs in the sequences that are radically distant from germlines and with examined examples we were unable to determine which tool is more correct or less wrong.

While renumbering CDRs, ANARCI reorganizes residues from the loop, with two flanking positions on each side. IMGT definitions of CDR boundaries are used for this regardless of the selected numbering scheme. In contrast to this RIOT renumbers CDR residues without flanking positions, from the target scheme loop definition. In edge case scenarios this leads to different region assignments of the residues. For example, with the absence of highly conserved tryptophan on IMGT 118, with a deletion on input sequence according to query-germline alignment, RIOT will show missing residue and FWR4 will start on position 119, where ANARCI smoothing logic will “pull” the nearest residue from CDR and place it at position 118. Furthermore the smoothing logic applied by ANARCI in IMGT space which is translated further to the user-selected schemes can lead to issues in Kabat-like schemes.

### Kabat/Chothia/Martin - AbNum, ANARCI, AntPack and RIOT compared

Of all therapeutic sequences, there were ∼200 conflicts in numberings. The majority of conflicts could be attributed to differences in CDR numbering heuristics or AbNum assigning the last residues of light chains to 106A instead of 107 in contrast to the remaining tools.

AbNum has two distinct heuristics for CDR numbering in Kabat derived schemes. For example, if present, deletions in CDR3 are placed to the left of the scheme-defined position for insertions (as defined in Table 2). On the other hand deletions in CDR1 are placed to the right of the insertion position. Such an approach was introduced in AbNum as an arbitrary design decision inferred from observed CDR sequence lengths. Reliance on observed commonly incomplete data can lead to non-generalizable solutions (as in case of whole Kabat scheme) therefore RIOT unifies CDR numbering heuristics so that all CDRs are numbered in the same way - if present - deletions are placed to the left of scheme-defined indel position. We believe such an approach improves the interpretability of the results.

After ignoring differences created as a result of either of the above reasons, a number of conflicting numberings remained. Some of them could be attributed to the fact that in some rare cases, ANARCI truncates the middle of the input sequence for Kabat-like schemes. Furthermore, ANARCI fails to number the sequence when there is a deletion on the J gene. On the other hand, AbNum fails to number some of the sequences. Finally, as it was in the case of IMGT, sequences that differ drastically from germlines can have different numberings.

## Discussion

Here, we introduced RIOT, software for high-throughput annotation of nucleotide and amino acid sequences of immunoglobulins with comprehensive germline databases for humans and mice derived from OGRDB (W. Lees et al. 2020). Our evaluations show that RIOT outperforms the leading nucleotide-sequence annotation tool, IgBLAST in performance, and the amino-acid sequence annotation tool, ANARCI in both performance and germline assignment accuracy. Though not as fast as the current leading numbering solution, AntPack, it offers a much wider functionality and precise germline annotations being within the range of its speed.

In large-scale Next-Generation Sequencing (NGS) studies, processing speed is crucial due to the massive volume of data generated (Dudzic et al. 2024; Parkinson and Wang 2024) that then needs to be effectively analyzed (Chomicz et al. 2024). Efficient processing allows for quicker analysis, enabling researchers to handle larger datasets and accelerate the pace of scientific discovery. With a typical processing pipeline consisting of both IgBLAST and ANARCI, RIOT significantly enhances this aspect by offering a performance that is approximately 8 times faster, by virtue of using a single tool rather than both of them. This improvement in speed is particularly beneficial in large-scale studies, where the ability to rapidly process and analyze data can lead to faster insights into immune responses and disease mechanisms.

Accurate germline assignment is essential in immunoglobulin analysis as it impacts the reliability of research findings. The accuracy with which immunoglobulin sequences are mapped to their corresponding germline genes affects the interpretation of B cell development and thus affinity maturation. Precise germline assignment is crucial for understanding immune system behavior, developing effective vaccines, and designing therapies for autoimmune diseases and infections. Lineage tree reconstruction is crucial in the analysis of NGS outputs from Phage Display or immunization that are employed in antibody-based drug discovery (Nouri and Kleinstein 2018).

Processing speed of modern numbering tools is a result of using fast sequence aligners and reducing the number of alignments being performed. The fastest solution, AntPack, achieves its performance by doing just 3 alignments to consensus sequences (H,K,L), by compromising germline assignment accuracy. We advocate for assigning germlines through alignment as we think it renders most biologically accurate results which might be crucial applications such as germlining-based humanization or lineage tree reconstruction.

While RIOT offers significant advancements in processing speed and efficiency, it is important to recognize its potential limitations. RIOT may produce unexpected results when the genes of the query sequence significantly differ from the closest genes in the RIOT database. This discrepancy can lead to incorrect gene assignments, potentially affecting the accuracy of the analysis. Moreover, RIOT may encounter difficulties in cases of frame-shifting modifications. These modifications alter the translation of the J gene-derived sequence segment, making it challenging to accurately determine the boundaries of the framework region.

Though RIOT can offer improvements over IgBLAST and ANARCI for immunoglobulins, it does not address T-cell receptors like the two pieces of software. By making the software available, we hope that the findings related to immunoglobulins can also be applied to T-cell receptors, advancing the biological study of these molecules as well as their therapeutic applications.

Even the best annotation software will not perform well if the underlying germline database lacks sufficient quality. The ongoing effort to build and maintain comprehensive reference germline databases is critically important. (W. D. Lees et al. 2023; Collins et al. 2023). These databases are fundamental to the functionality of tools like RIOT, as they provide the essential reference points for sequence comparison and assignment. Continuous updates and expansions of these databases are crucial to keep up with the genetic diversity observed in different populations and to adapt to new immunological challenges (Peng et al. 2021). An effort that must not be forgotten is the serialization of database versions used for annotations.

In conclusion, RIOT introduces significant improvements in the analysis of immunoglobulin sequences, particularly in terms of processing speed and integration of functionalities. We believe RIOT will significantly aid in the analysis of immunoglobulins by providing a unified platform for comparing sequences against a reference germline database.

